# Minimizing marine ingredients in diets of farmed Atlantic salmon (*Salmo salar*): Effects on growth performance and muscle lipid and fatty acid composition

**DOI:** 10.1101/328716

**Authors:** Maryam Beheshti Foroutani, Christopher C. Parrish, Jeanette Wells, Richard Taylor, Matthew Rise, Fereidoon Shahidi

**Author notes:** Corresponding author: (CP).

## Abstract

Due to limited fish meal and fish oil resources and their high costs for the aquaculture industry, it is necessary to find alternative sustainable sources of protein and lipids. Therefore, seven different diets were formulated with different protein and lipid sources to feed farmed Atlantic salmon, and their effects on growth performance, muscle lipid class, and fatty acid composition were examined. Growth performance indicated that the diet with the lowest fish meal and fish oil content resulted in the lowest weight gain and final weight, followed by the diet containing the highest level of animal by-products. The lipid class analysis showed no statistical difference in the muscle total lipid content using different diets. However, significant statistical differences were observed among the main lipid classes; triacylglycerols, phospholipids, and sterols. The diet containing 1.4% omega-3 long-chain fatty acids resulted in the highest content of triacylglycerols and phospholipids. Diets containing medium and low levels of fish oil and fish meal, respectively, led to as high a level of ω3 fatty acids in muscle as when fish were fed diets with high levels of fish meal and fish oil. The results of this study suggest that feeding a diet containing low levels of fish meal and moderate levels of fish oil does not significantly affect ω3 fatty acid composition in muscle. Fish meal could be reduced to 5% without affecting growth as long as there was a minimum of 5% fish oil, and animal by-products did not exceed 26% of the diet.

## Introduction

Fish meal and fish oil have been the most used ingredients in aquafeed manufacturing as they are excellent sources of proteins and lipids. From 1995 to 2004, the global consumption of fish meal and fish oil approximately doubled [1]. However, the production of fish meal has not changed significantly in the past 30 years, and fish oil has been produced at much lower levels [2]. Fish meal contains valuable protein with high digestibility as well as essential vitamins and minerals [3]. Fish oil is a valuable primary source of lipid providing the required and beneficial omega (ω)-3 polyunsaturated fatty acids (PUFA). Aquaculture’s high reliance on fish oil has recently made researchers study alternative lipid sources to reduce the pressure on the consumption of fish oil [4]. However, finding sustainable ingredients that could provide the required nutrition for fish have been found to be difficult as these alternative sources are not as nutritious as fish oil [5].

Marine fish and salmonids require arachidonic acid (ARA, 20:4ω6) and high levels of essential omega-3 fatty acids, eicosapentaenoic acid (EPA, 20:5ω3) and docosahexaenoic acid (DHA, 22:6ω3), as they cannot biosynthesize them easily. However, fish have some ability to supply these essential fatty acids by synthesizing them from shorter chain PUFAs (e.g. 18:3ω3 and 18:2ω6). Therefore, it is important that high levels of PUFAs are available in the diet [6].

Fish are a great source of ω3 in the human diet. EPA and DHA play an important role in different functions, including nervous system, photoreception, reproduction, as well as in reducing the risk of cardiovascular and inflammatory diseases [7]. Therefore, it is important to ensure that these essential fatty acids are provided in fish diets as they contribute to the health of fish and the human consumer. Growth, flesh quality, and health of fish are improved as the level of EPA and DHA in the diet increases [6].

Proteins constitute the major part of the organic material in fish tissues (70% of dry weight). Hence, protein content and quality in fish diets play a very important role in producing a high quality fish and growth performance of fish is directly dependent on the level of protein in the diet as it provides essential amino acids. However, minimizing the usage of protein is important as it is one of the most expensive components of the feeds.

The increasing need of fish meal and fish oil has resulted in higher prices of these sources. High costs and limited availability of fish meal and fish oil have led to several evaluations of the effect of their replacement with alternative sustainable sources, such as those derived from terrestrial plants and animals, in the diets of different fish species. While partial replacement generally was successful in terms of growth performance, full replacement mostly resulted in lower content of ω3 and essential fatty acids, as well as poor growth performance [8], [9], [10], [11].

Several studies have evaluated the effect of replacing fish meal and fish oil with terrestrial plant sources in feeds for farmed animals [12] as they are sustainable sources of energy for fish growth that are readily available at lower prices [13], [14]. However, one problem that limits full replacement of fish oil with plant oils is the low level of ω3 fatty acids and high levels of ω6 and ω9 fatty acids. Plant oils can provide the same relative ratios of saturated fatty acids, monounsaturated fatty acids (MUFA) and PUFA as found in fish oil, but not the level of highly unsaturated fatty acids in fish oil [15], [16]. Nonetheless plant oils can replace a major amount of fish oil in the diets [8], [9], [10], [11] with no significant influence on fish growth as long as sufficient levels of essential fatty acids are provided [12]. To minimize any reduction in growth rate, and nutritional quality in terms of the health benefits of farmed fish to human consumers, potential substitutes for fish oil should avoid excessive deposition of 18:2ω6, retain high levels of ω3 HUFA and provide sufficient energy in the form of saturated and MUFAs [17]. It is claimed that as long as essential fatty acid requirements are met, fish oil can be substantially replaced by MUFA-rich alternative sources. Rapeseed oil is one of the best candidates to replace fish oil because it contains a high portion of MUFA, varying between 55 and 72% of total fatty acids [11]. MUFAs cannot be bioconverted to essential fatty acids like LA, ALA, and ω3 long-chain PUFAs, so the main reason for using plant oils rich in MUFAs is their ability to provide the required energy [11].

The effect of replacing fish oil with rapeseed oil in diets of Atlantic salmon (*Salmo salar*) on tissue lipid composition was evaluated in [13]. The results showed no effect on growth performance, feed efficiency, in the samples of liver, muscle. Fish oil diets resulted in high concentrations of lipids in the muscle. However, high lipid levels occurred in the liver in fish fed 100% rapeseed oil. The result of muscle lipid fatty acid composition analysis revealed that 18:1ω9, 18:2ω6, and 18:3ω3 all increased with increasing replacement level. Increase of rapeseed oil in the diet, however, considerably reduced total saturated fatty acids and 20:5ω3 and 22:6ω3. It was suggested that rapeseed oil is a great candidate for replacing fish oil in the diet of Atlantic salmon [13. However, replacement at 50% or higher significantly reduces the concentrations of 20:5ω3, 22:6ω3, and the ratio of ω3 to ω6 in the muscle making it less beneficial for humans.

Another alternative source to replace fish meal is terrestrial plant proteins. Fish meal and fish oil were replaced [9] with a blend of plant proteins and vegetable oils to make a sustainable diet for Atlantic salmon, meeting the nutritional requirements. The result of their study suggested four times greater efficiency in terms of fish meal consumption at 80% replacement level with plant proteins.

Rendered terrestrial animal products can also be used as an alternative protein source to fish meal [18], [19]. One of the animal protein sources widely used in fish diets is animal by-products, given that carnivorous species prefer replacements of animal origin rather than plant sources due to palatability. The use of animal by-products have several advantages, including being free from antinutritional factors such as phytic acid, phosphorus, or indigestible complex carbohydrates. Furthermore, they contain low amounts of carbohydrates, and are high in crude protein and crude lipids, as well as vitamins such as B_12_, and trace minerals such as iron, cobalt, and selenium [20]. In addition to the good nutritional value of poultry by-product meal and hydrolyzed feather meal, they have a very competitive cost advantage over fish meal [21].

While there have been several studies on replacing marine ingredients with alternative sustainable sources, more effective replacement levels are required. Therefore, the objective of this study was to evaluate how using alternative diets with low content of marine resources can affect growth performance, and muscle lipid class and fatty acid composition in farmed Atlantic salmon. This study improves the understanding of the nutrition requirements of Atlantic salmon under various dietary conditions, and provides an insight into the potential performance of future aquafeed alternatives.

## Materials and methods

### Experimental diets

Seven different diets were produced by EWOS Innovation AS in Norway. The diets were formulated using different levels of fish meal, fish oil, animal by-products, vegetable oil, and vegetable protein, yielding different levels of DHA+EPA. The seven diets were characterized according to the most critical component used and designated as marine, with high levels of fish meal and fish oil; medium marine, containing medium levels of fish meal and fish oil; animal by-product, composed of a high proportion of animal by-products; vegetable protein, including a high level of vegetable protein; ω3LC0, which contained approximately 0% long-chain ω3 fatty acids (LC ω3 FAs); ω3LC1, with 1% of LC ω3 FAs; and ω3LC1.41, with 1.4% of LC ω3 FAs (Table 1).

**Table 1.**
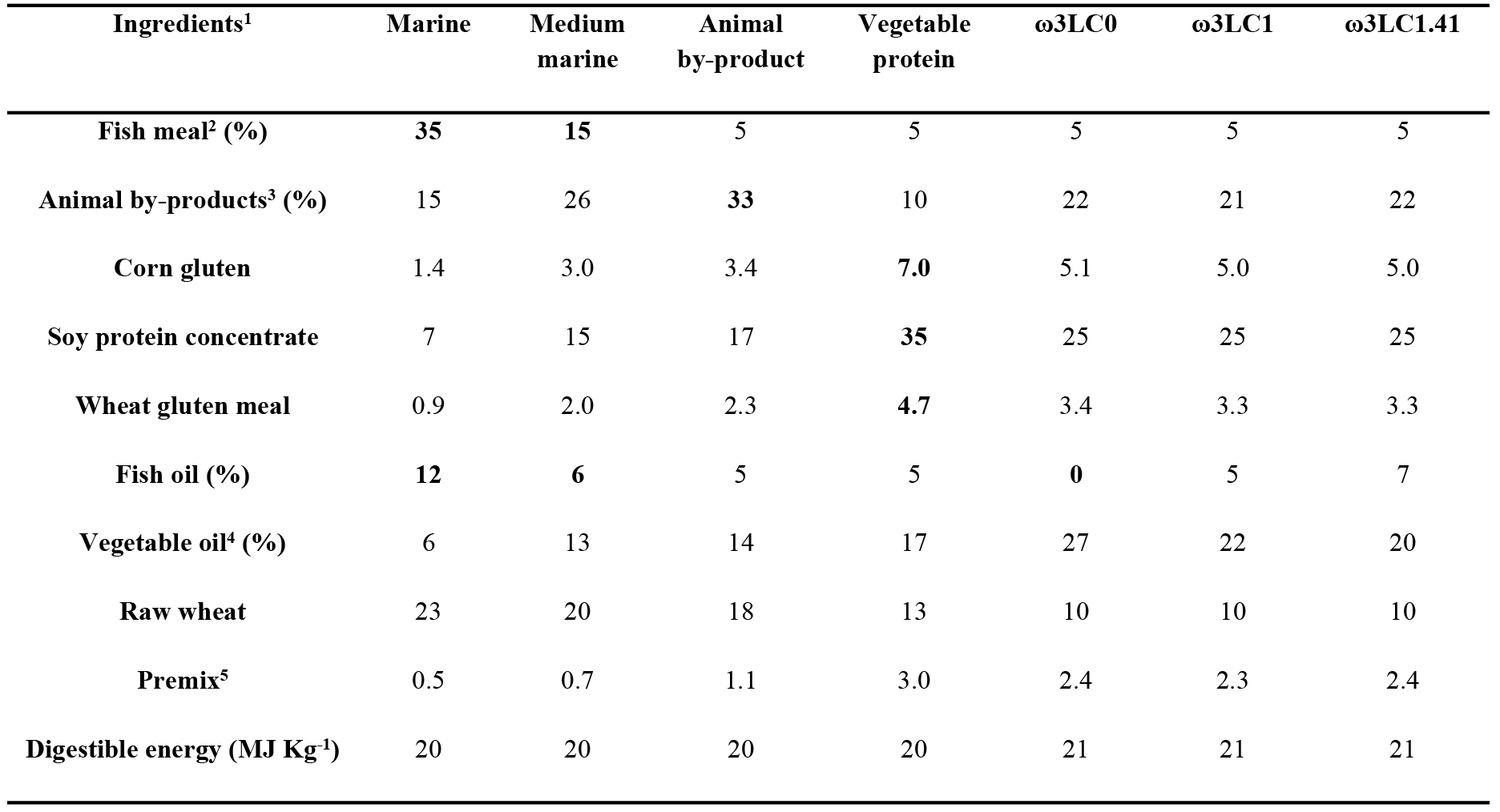
Experimental diet ingredients.

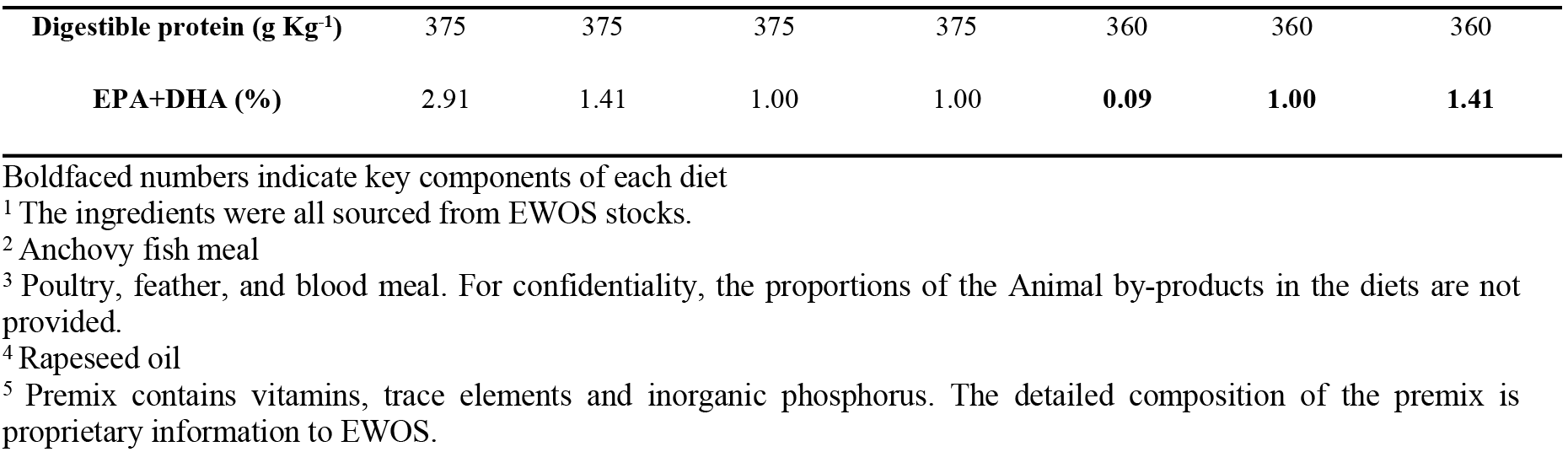

### Experimental fish and feeding

Atlantic salmon (*Salmo salar*) smolts were obtained from Northern Harvest Sea Farms in Stephenville, NL, Canada. Fish were tagged with passive transponders (PIT) and maintained at 11±1°C in a flow-through seawater system under a 12-h light photoperiod in the Dr. Joe Brown Aquatic Research Building in St. John’s, Newfoundland, Canada. Fish (1148 at 139-232 g each) were distributed randomly in 28 tanks (620 L, 4 tanks per diet). There were 41 fish per tank until day 0, when one fish/tank was sampled. Fish were hand fed 5 mm experimental pellets to satiation twice a day for 14 weeks. Feed consumption was measured weekly and fish weight and fork length were recorded at the beginning and end of the experiment.

### Tissue sampling

Ten fish were randomly sampled at week 0, followed by 5 fish per tank at week 7 and 14. Fish were euthanized with an overdose of MS-222 (400 mg L^−1^, Syndel Laboratories, Vancouver, BC, Canada), then fork length and weight were measured. Gut and liver were removed, weighed, sampled, and placed on aluminum weigh boats. Norwegian quality cuts (NQC) (area from directly behind the dorsal fin to the anus) were removed from the fish body and weighed. With the dorsal fin cut to the right, the skin was cut along the dorsal side, peeled back to expose the skeletal white muscle (hereafter referred to as muscle), and a strip (approximately 0.5 g) of muscle was removed and placed into lipid clean test tubes (rinsed with methanol and chloroform three times each) for lipid analysis. The tubes had been weighed after ashing at 450°C for 8 hrs, and the Teflon lined caps were rinsed three times with methanol and chloroform. Vials were kept on ice during sampling. After sampling, 2 ml of chloroform was poured on the tissues in the tubes and the remaining space was filled with nitrogen. The tubes were then sealed and stored at −20°C. All procedures, including handling, treatment, euthanasia, and dissection were performed according to the guidelines from the Canadian Council of Animal Care (approved Memorial University Institutional Animal Care Protocol 14e71-MR).

### Lipid extraction and lipid class determination

Samples were ground in methanol:chloroform:water and extracts were evaporated to volume under a gentle stream of nitrogen before sealing and storing at −20°C [22]. Lipid class composition was determined through a three-step development method using thin-layer chromatography-flame ionization detection [23]. Lipid extracts and standards were spotted on Chromarods and the rods were developed and then scanned in an Iatroscan. Data were collected using PeakSimple software.

### Fatty acid methyl ester (FAME) derivatization

Fifty microlitres of lipid extract was transferred into 15 ml lipid clean vials and concentrated to dryness. Methylene chloride (1.5 ml) and 3 ml of Hilditch reagent (1.5 H_2_SO_4_: 98.5 anhydrous methanol) were added, then vials were vortexed and sonicated for 4 min. They were filled with nitrogen, capped and heated at 100°C for 1 hr. Saturated sodium bicarbonate solution (0.5 ml) and 1.5 ml hexane were added to the samples which were vortexed, followed by removing the upper, organic layer. The vials were then dried and refilled with hexane to approximately 0.5 ml. They were filled with nitrogen, capped, sealed with Teflon tape, and finally sonicated for another 4 min to re-suspend the fatty acids.

### Lipid oxidation

Lipid oxidation was measured according to the thiobarbituric acid reactive substance (TBARS) method [24]. To measure lipid oxidation, 1 gram of muscle sample (two replicates per fish) was weighed and transferred into centrifuge tubes (plus 1 blank). The sample was homogenized with 5 ml of 5% (w/v) trichloroacetic acid (TCA) by polytron. Samples were centrifuged at 3000 rpm for 10 min and the supernatant (top layer) was filtered through a 0.45 μm pore syringe filter. Five ml of 0.08 M thiobarbituric acid (TBA) and 2.5 ml of TCA were added and heated in a boiling water bath at 94±1°C for 45 min then cooled to room temperature. Finally, the absorbance was measured at 532 nm using a UV-spectrophotometer. Thiobarbituric acid reactive substances (TBARS) values were calculated using a standard curve. Standard curves were prepared using 1,1,3,3-tetramethoxypropane as a precursor of malondialdehyde (MDA; 0 - 10 ppm).

### Statistics

To ensure representative fish were sampled for analysis, only fish with weight gains within the range of twice the standard deviation from the overall tank weight gain means were considered. Correlation and regression analyses were conducted using Minitab version 17 to compare diet ingredients, lipid and fatty acid composition of muscle and diet, and growth characteristics. For statistical analysis of growth data, lipid class, and fatty acid data, nested general linear models were combined with Tukey pairwise comparisons using Minitab to determine the difference between tanks and diets. The normality of residuals was evaluated with the Anderson-Darling normality test. If the test failed (p<0.05), a one-way analysis of variance (ANOVA) on ranks was performed in SigmaPlot version 13.

## Results

### Experimental diet composition

Moisture levels in the feeds ranged from 3.88 to 6.39% and the major lipid class was triacylglycerol ranging from 92 mg g^−1^ in the animal by-product diet to 173 mg g^−1^ in the ω3LC0 diet (Table 2). All diets had the same concentrations of 16:0, ΣSFA, and ΣPUFA as well as the same ratios of PUFA/SFA and DHA/EPA. The three diets with 5% fish meal and ~22% animal by-products, ω3LC0, ω3LC1, and ω3LC1.41 had statistically the same content of 18:1ω9, 18:2ω6, 18:3ω3, and 20:4ω6. The 20:5ω3 and 22:6ω3 contents were significantly lower in the ω3LC0 diet than in the marine, medium marine and ω3LC1.41 diets (Table 3).

**Table 2.** Lipid classes in experimental diets, as fed (mg g^−1^ wet weight)

| Lipid classes | Marine | Medium marine | Animal by-product | Vegetable protein | ω3LC0 | ω3LC1 | ω3LC1.41 |
| --- | --- | --- | --- | --- | --- | --- | --- |
| (mg g<sup>-1</sup> wet weight) | n=6 | n=6 | n=9 | n=6 | n=9 | n=9 | n=9 |
| **Total lipid** | 161.1±64.5<sup>ab</sup> | 173.4±46.7<sup>ab</sup> | 140.1±39.6<sup>b</sup> | 159.3±27.6<sup>ab</sup> | 229.2±59.2<sup>a</sup> | 203.5±25.6<sup>ab</sup> | 208.7±38.7<sup>a</sup> |
| **Triacylglycerol** | 93.3±45.2<sup>b</sup> | 96.1±28.1<sup>b</sup> | 92.1±40.6<sup>b</sup> | 115.4±27.3<sup>ab</sup> | 173.2±55.0<sup>a</sup> | 149.6±27.7<sup>ab</sup> | 147.8±34.9<sup>ab</sup> |

Data are mean ± standard deviation for n replicates

**Table 3.**
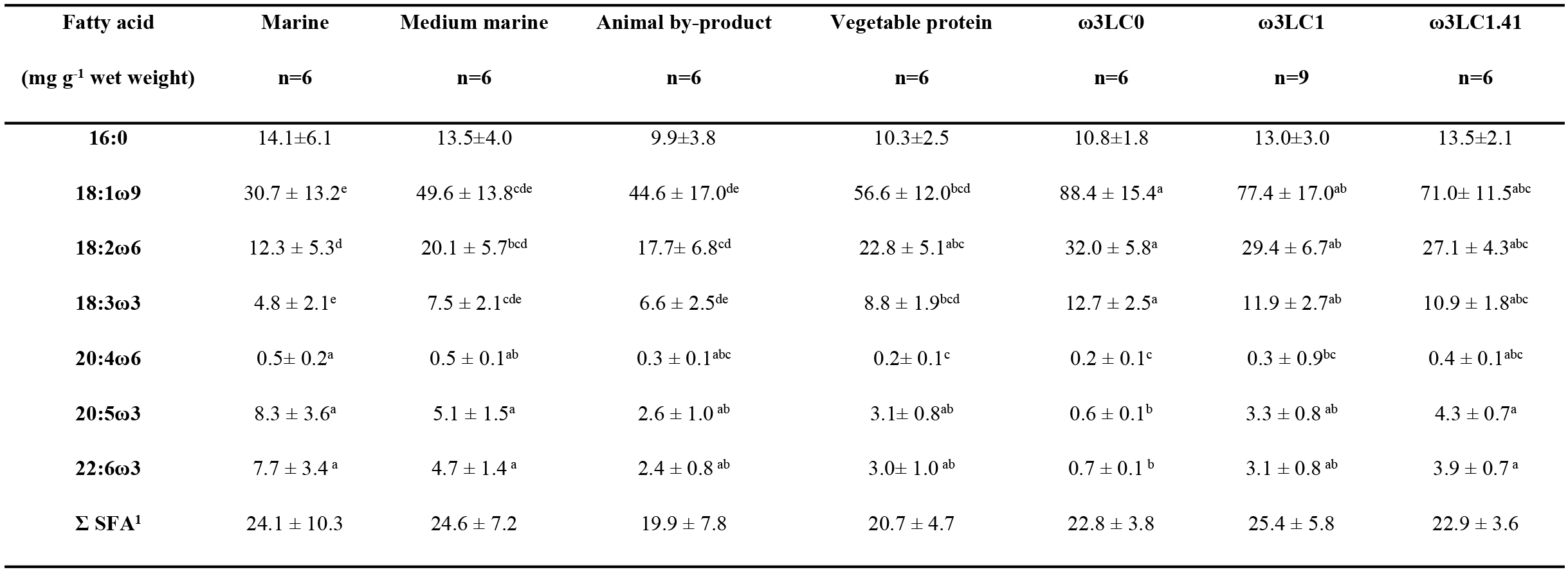
Fatty acids composition of experimental diets, as fed (mg g^−1^ wet weight)

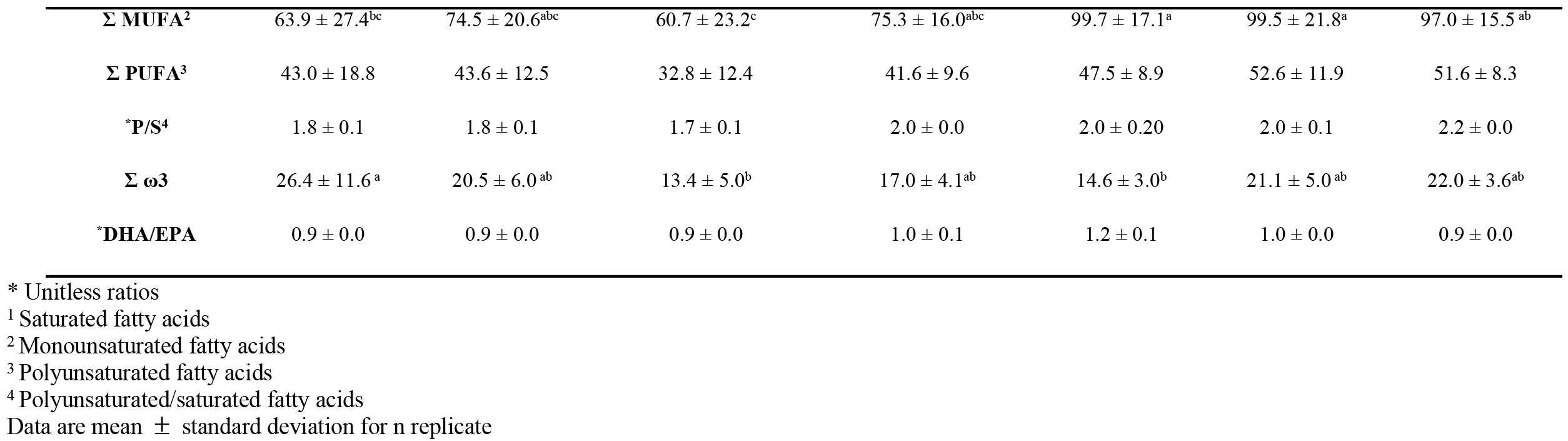

### Growth performance

After 14 weeks of feeding, final weights were lowest when using the diet with the lowest fish meal and fish oil content (ω3LC0), followed by the diet containing the highest level of animal by-products. Fish fed diets ω3LC0, ω3LC1, and ω3LC1.41 had the lowest hepatosomatic index (HSI) and fish fed the marine diet had the highest HSI (Table 4). The lowest specific growth rate (SGR) was obtained with the ω3LC0 and animal by-product diets (Table 4). The condition factor (CF) was not influenced due to relatively similar growth in body weight and length, which shows that the diets were well designed.

**Table 4.**
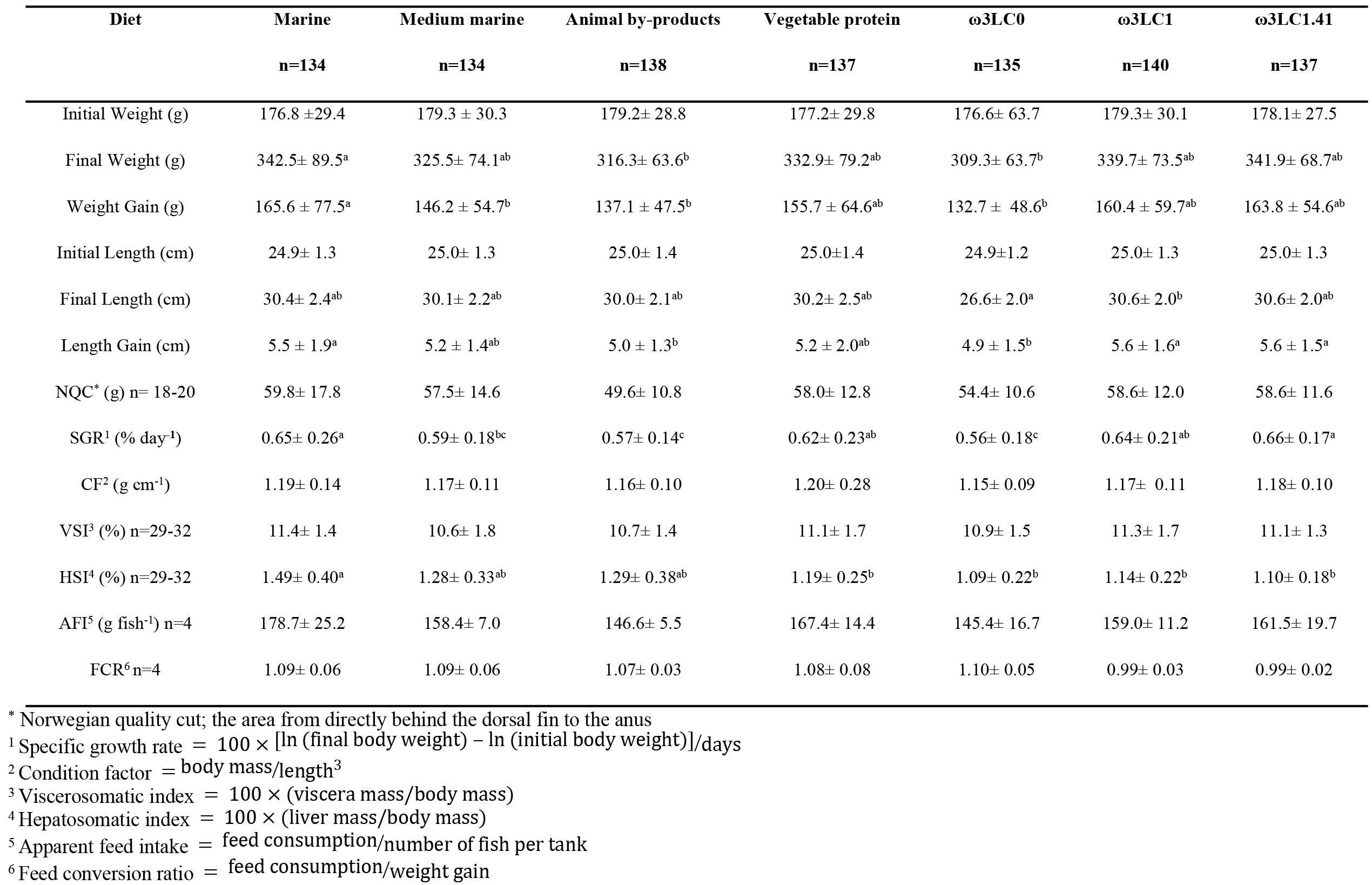
Salmon growth data - week 14.

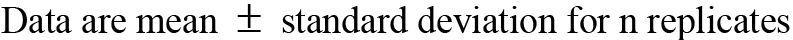

### Muscle lipid class composition and fatty acid composition

There was no significant difference in total lipids among the different treatments but muscle triacylglycerol was lowest with the animal by-product diet and highest with the ω3LC1.41 diet (Table 5). Fish fed the marine and animal by-products diets showed the highest and lowest level of sterol in muscle tissues respectively (Table 5). Diet ω3LC0 resulted in significantly higher contents of 20:4ω6 than medium marine diet. Animal by-products, ω3LC0, and ω3LC1.41 diets led to the highest DHA/EPA ratio, while the marine diet resulted in the lowest ratio (Table 6).

**Table 5.** Lipid classes of muscle tissue before and after 14 weeks of feeding (mg g^−1^ wet weight)

| Lipid classes | Initial | Marine | Medium marine | Animal by-products | Vegetable protein | ω3LC0 | ω3LC1 | ω3LC1.41 |
| --- | --- | --- | --- | --- | --- | --- | --- | --- |
| (mg g<sup>-1</sup> wet weight) | n=10 | n=16-19 | n=16-19 | n=16-19 | n=16-19 | n=16-19 | n=16-19 | n=16-19 |
| **Total lipid** | 15.2±7.6 | 17.8±14.0 | 9.3±5.2 | 8.6±4.7 | 13.7±5.5 | 10.6±6.0 | 17.2±12.3 | 18.1±15.3 |
| **Triacylglycerol** | 7.2±4.8 | 8.6±7.7<sup>ab</sup> | 5.6±4.5<sup>ab</sup> | 3.4±2.1<sup>b</sup> | 7.9±4.9<sup>ab</sup> | 5.1±3.4<sup>ab</sup> | 8.3±6.4<sup>ab</sup> | 9.2±6.7<sup>a</sup> |
| **Sterol** | 0.26±0.17 | 0.87±0.77<sup>a</sup> | 0.17±0.09<sup>ab</sup> | 0.20±0.33<sup>b</sup> | 0.23±0.04<sup>a</sup> | 0.26±0.16<sup>a</sup> | 0.29±0.27<sup>ab</sup> | 0.28±0.18<sup>a</sup> |
| **Phospholipid** | 5.0±2.4 | <u>2.8±0.8<sup>b</sup></u> | 2.9±1.3<sup>b</sup> | 3.9±1.5<sup>ab</sup> | 3.8±0.6<sup>ab</sup> | 4.2±2.8<sup>ab</sup> | 4.4±2.5<sup>ab</sup> | 5.2±3.2<sup>a</sup> |

Underlined data represent values that are statistically different from week 0
Data are mean ± standard deviation for n replicates

**Table 6.**
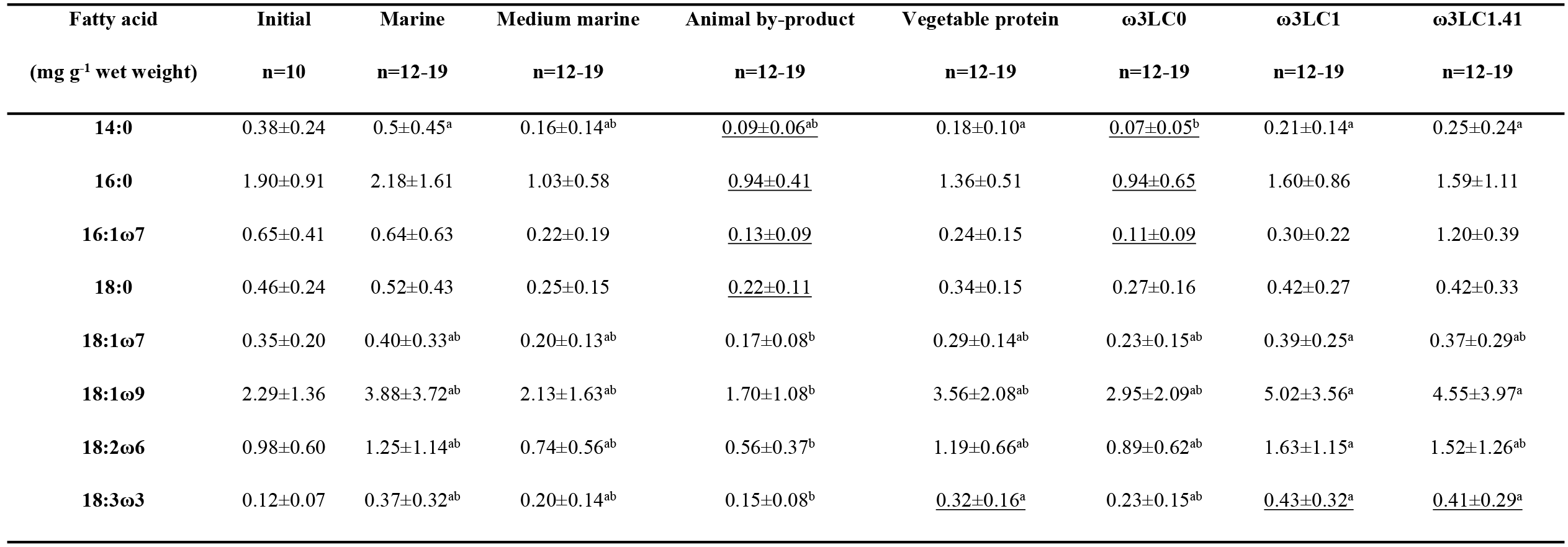
Fatty acid composition of muscle tissue before and after 14 weeks of feeding (mg g^−1^ wet weight)

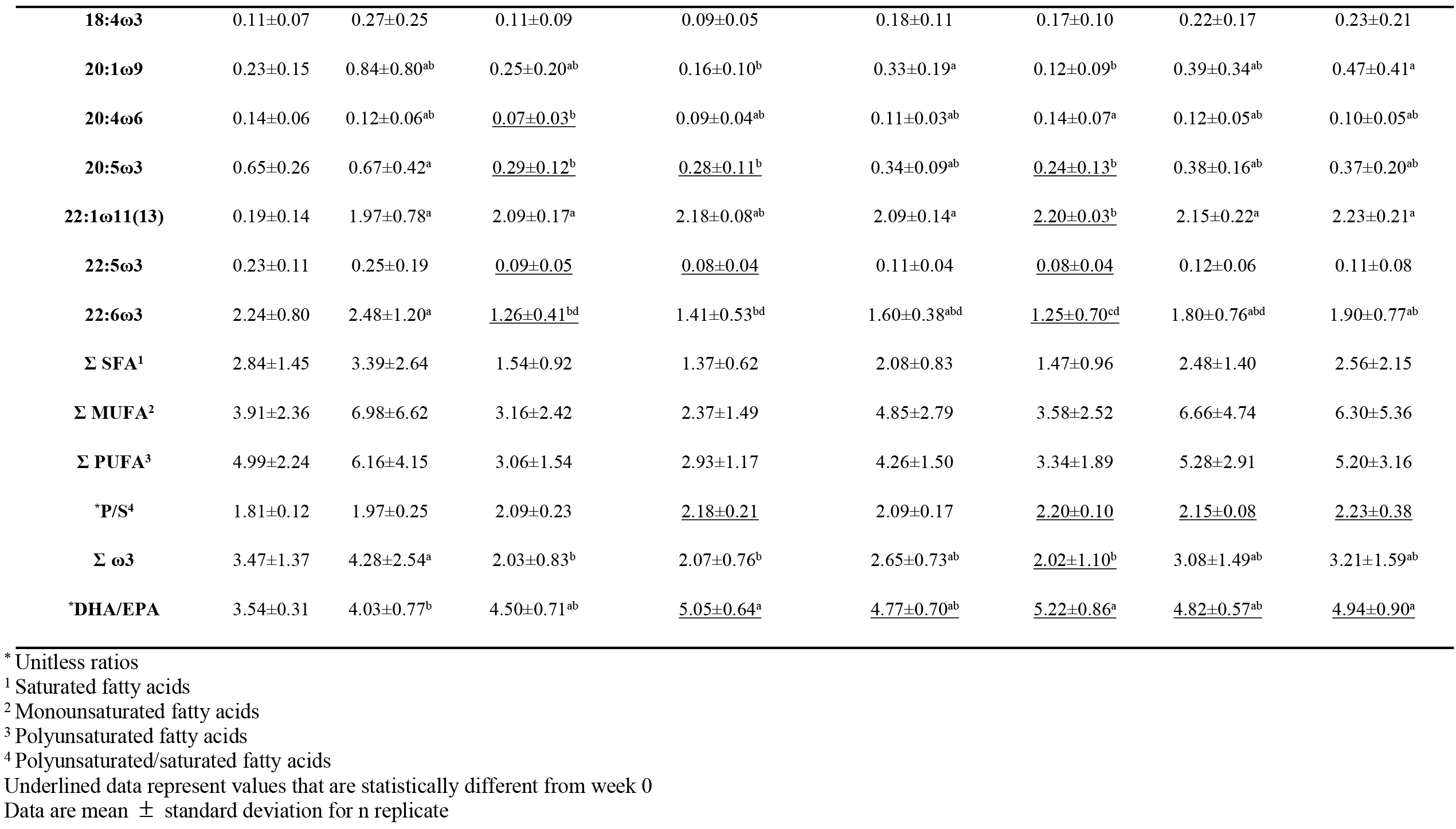

### Correlation analysis

Apparent feed intake, final weight and weight gain showed a significant positive correlation with the amount of fish oil in the diet, as did HSI; however, SGR, final length, NQC, VSI and CF did not correlate significantly with fish oil level. Σω3 in the diet and muscle had positive correlations with growth performance characteristics (r=0.82-0.87, Table S1) but there were significant positive correlations between HSI and fish meal, fish oil, and EPA+DHA (r=0.77-0.88). By contrast, there was an inverse correlation between HSI and vegetable oil content in the diet (−0.942, P<0.01), which is why the diet with the highest inclusion of vegetable oil (ω3LC0) resulted in the lowest HSI (Fig. S1, Table 4). Animal by-products inclusion showed a negative correlation with condition factor (CF, Table S1), while there was a positive linear relationship between the content of 22:6ω3 in diet and muscle dietary essential fatty acids and muscle fatty acids for 20:5ω3 and 22:6ω3 (P<0.05, R^2^ = 75.8% and P<0.05, R^2^ = 60%, respectively, Table S2 and Fig. S2).

### Lipid oxidation

Lipid oxidation was also studied to measure the extent it could be affected by muscle tissue lipids, as high content of PUFA could make them susceptible to oxidation. Oxidation results were represented by TBARS values for marine, animal by-product, and ro 3LC1.41 diets with mean values of 1.15±0.61, 1.52±0.65, and 1.01±0.59 mg MDA kg^−1^ of sample (mg of malondialdehyde equivalents per kg of muscle sample), respectively.

## Discussion

While the global consumption of marine ingredients by aquaculture continues to increase, their production has not changed significantly [2] and has even declined slightly in recent years [25]. Consequently this study evaluated the effects of replacing fish meal and fish oil in aquafeeds with soy protein concentrate, poultry, feather, and blood meal, and with rapeseed oil.

Cultured fish need lipid, protein, energy, vitamins and minerals in their diet for growth, reproduction, and other normal physiological functions. Lipid contributes to the structure of biomembranes [26] and has a critical role in providing energy for animal tissues and is the source of essential fatty acids. The poor performance of fish fed the ω3LC0 diet is undoubtedly associated with the complete lack of fish oil in this diet [27]. Fish meal contains valuable eicosapentaenoic acid (EPA, 20:5ω3) and docosahexaenoic acid (DHA, 22:6ω3) and a relatively high content of fish meal in the diet reduces the possibility of having low levels of essential fatty acids [12]. The content of fish meal in the diet of Atlantic salmon was reduced to 25% in [10] with no significant change in growth and feed conversion. However, the present study further reduced the fish meal content to as low as 5%.

The ω3LC0 diet also resulted in the lowest HSI which is contrary to the results of other studies on salmonids (e.g. [28], [29]), in which HSI was statistically un affected by the increase of vegetable oil content in the diet. However in those studies the maximum vegetable oil content was 20%, whereas here the ω3LC0 diet had the highest inclusion of vegetable oil at 27%.

Correlation analysis showed that growth performance characteristics are positively correlated with total concentrations of ω3 fatty acids in the diets, but interestingly there was no significant correlation with individual ω3 fatty acids in the diets suggesting that an improvement in growth performance may be obtained irrespective of ω3 fatty acid chain-length or degree of unsaturation. In turn this indicates an interchangeability among these fatty acids in terms of function or biochemically through modification of chemical structures, e.g. chain elongation and desaturation.

With the exception of the animal by-product diet, diets containing ≥5% fish meal and ≥5% fish oil gave the same final weight, and length gain, probably because the minimum level of essential fatty acids was provided [30]. However, as with the ω3LC0 diet, the animal by-product diet resulted in significantly lower growth despite containing 5% fish meal and 5% fish oil, which may relate to lipid oxidation. While muscle in fish fed the animal by-product diet was expected to be less exposed to lipid oxidation due to the lower content of PUFAs (Table 6), the results revealed that these fish muscle samples were oxidized significantly more than the ones fed ω3LC1.41. This may reflect the types of materials used for animal by-products and how they were handled. Nonetheless, the oxidation results suggested that lipid rancidity was not high in any of the three diet treatments studied, as the TBARS values were significantly lower than the results of other studies on salmon (e.g. [31]). A TBARS value of about 16.5 mg MDA kg^−1^ was obtained in [31] after 6 days, while the TBARS values in this study did not exceed 3.13 mg MDA kg^−1^ (mean of 1.52 mg MDA kg^−1^). Also, lower levels of animal by-products may be well tolerated. It was shown in [32] that 7% inclusion of porcine blood meal had no negative effects on salmon performance compared with fish meal based control diets. The animal by-product diet also resulted in the lowest concentrations of muscle triacylglycerols which is consistent with the down-regulation of fatty acid synthesis in the livers of these fish [33].

The vegetable protein and ω3LC1 diets were remarkable in giving the same results for final weight, weight gain, final length, length gain, and SGR as when the marine diet was used, despite having less than half the amount of fish oil in the diet and as little as 1/7 of the fish meal. In addition, the vegetable protein diet was found to increase the immune response of these fish [34].

For fish fed diets containing 5 – 7% fish oil and 5-15% fish meal, there was no significant difference in the concentrations of DHA nor EPA in the muscle. EPA and DHA have anti-inflammatory effects, which make fish resistant to diseases [1]. As expected, DHA and EPA was lowest with the ω3LC0 diet, but ARA was highest (Table 6). This despite the concentration of ARA being lowest in this diet along with the vegetable protein diet. This diet contains a high inclusion of rapeseed oil which has a high level of 18:2ω6, so conversion to 20:4ω6 [35] in muscle tissues is strongly indicated.

While five of the treatments gave no significant difference in muscle DHA and EPA concentrations the two with the lowest and highest levels of fish oil did give significant differences, as expected ([36], [15], [37]). This resulted in significant regressions between 20:5ω3 and 22:6ω3 in diets and muscle tissues. The slope of the regression line for dietary content of 20:5ω3 and 22:6ω3 against their contents in muscle tissue indicates the extent of this dependency. The higher slope for 22:6ω3 (0.30 mg g^−1^ wet weight) compared to 20:5ω3 (0.08 mg g^−1^ wet weight) showed that 22:6ω3 was deposited four times more than 20:5ω3 in muscle tissue. A similar result was observed when fish oil was replaced with crude palm oil in Atlantic salmon diets [17], where 22:6ω3 (slope = 0.81) was more deposited than 20:5ω3 (slope = 0.58) in muscle tissues when proportions (% total fatty acids) were compared. Similar to other studies (e.g. [13], [36], [28], [9], [38], [39]), the concentration of long chain MUFAs (20:1ω9 and 22:1ω11(13)) in the muscle tissues were highest for fish fed marine based diets. However, unlike the previous studies in which the replacement of marine based ingredients generally resulted in lower levels of long chain MUFAs, the use of other diets led to as high levels of 20:1ω9 and 22:1ω11(13) in muscle as when marine based diets were fed (Table 6).

The fatty acid compositions of muscle tissues (Table 6) show that the sum of EPA and DHA levels ranges from 1.49 to 3.15 mg g^−1^ wet weight. According to the Canada’s Food Guide [40], one serving of fish is 75 g. Therefore, a serving of Atlantic salmon fed the seven dietary treatments includes EPA+DHA levels ranging from 112 to 236 mg which would be less than the daily requirement (250 mg) recommended by the World Health Organization [41], even for fish fed the marine diet.

## Conclusion

This study evaluated the effects of minimizing marine resource utilization in diets of farmed Atlantic salmon on growth and muscle lipid composition. By replacing fish meal and fish oil in aquafeeds with animal by-products and rapeseed oil at levels used in this study (33 and 27%, respectively), marine resource utilization was reduced to 10% or less. This affected growth, and lipid class and fatty acid composition of muscle tissue, unlike with lower replacement levels of 26% and 22%, respectively.

## Acknowledgements

The authors would like to acknowledge Danny Boyce and the Joe Brown Aquaculture Research Building staff for their valuable help with fish rearing and sampling.

## Supporting information

**Table S1. Correlation analysis r values among diet ingredients, growth performance, diet (D) lipid classes, diet fatty acid composition, muscle (M) lipid classes, and muscle fatty acid composition (data with*, **, and *** represent P≤0.05, P ≤0.01, and P≤0.001, respectively).**

**Table S2. Correlation analysis r values among diet ingredients, diet lipid classes, diet (D) fatty acid composition, muscle (M) lipid classes, and muscle fatty acid composition (data with*, **, and *** represent P≤0.05, P ≤0.01, and P≤0.001, respectively).**

**Figure S1. Regression analysis between vegetable oil percentage in the diet and hepatosomatic index.**

**Figure S2. Regression analyses between amount of 20:5ω3 (a) and 22:6ω3 (b) in the diet and muscle tissue.**

